# A versatile bulk electrotransfection protocol for mouse embryonic fibroblast and iPS cells

**DOI:** 10.1101/763615

**Authors:** Shahin Eghbalsaied, Iqbal Hyder, Wilfried A. Kues

## Abstract

A square-wave pulsing protocol was developed using OptiMEM-GlutaMAX for high efficient transfection of mouse embryonic fibroblast (MEF) and induced pluripotency stem (iPS) cells. An electrotransfection efficiency of > 95% was repeated for both MEF and iPS cells using reporter-encoding plasmids. The protocol was very efficient for plasmid size ranging from 6.2 to 13.5 kb. A high rate of targeted gene knockout (> 95 %) was produced in Venus transgenic cells using indels formation. Targeted deletions in the Venus transgene were performed by co-electroporation of two gRNA-encoding plasmids. In conclusion, this plasmid electrotransfection protocol is straight-forward, cost-effective, and efficient for CRISPRing mouse primary cells.

## Background

The CRISPR (clustered regularly interspaced short palindromic repeats)–Cas9 (CRISPR-associated 9) protein nuclease system is a straight-forward, versatile, highly efficient, and nascent tool for genome editing of various organisms. Using ‘all-in-one’ expression vectors containing expression cassettes for guide RNAs and Cas9 nuclease/nickase it is possible to conduct CRISPR/Cas9 studies through a low-cost and straight-forward approach (1). However, the efficiency of gene editing using plasmid-based delivery methods remains relatively low, which subsequently increased attentions toward using alternate methods of employing the Cas9 protein and/or guideRNAs and using ribonucleoproteins (RNPs) (2, 3).

Basically, gene transfer into cells can be achieved via electroporation, lipofection, or viral transduction. Although viral gene transfer is very efficient, it requires time, skilled staff, and high levels of safety issues, whereas it has a limitation in the insert size and more importantly enhanced risks for clinical researches (4). On the other side, lipofection suffers from a low efficiency. Electroporation is an approach to instantly create several pores in the cell membrane using a burst electric pulse and to mediate the transfer of micro- and macro-molecules into cells, embryos, tissues, and organs (5). From the early papers on electroporation (6), it has been evident that gene transfer via electroporation is simple, easily applicable, and also efficient compared to lipofection (6). Although electroporation has been widely accepted as the main gene transfer tool in various laboratories around the world, the underlying mechanism has not been completely understood (7). Therefore, optimization of various factors, such as the electroporation medium, cuvette type (path width, length, and high), as well as the pulsing method which includes the amount, number, duration, and interval of pulses, is needed to have a high electrotransfection efficiency particularly in primary cells. Based on the above-mentioned electroporation parameters various types of electroporation-based devices, such as nucleofection (8), nano-electroporation (9), mechanical-electrical approach (7), and microfluidic membrane deformation (2) as well as new generation of electroporator devices (10), have been invented. However, still there is a huge room to improve the electroporation efficiency to transfer exogenous DNAs into cells, specifically for large plasmids (7, 11).

Here, we introduced a highly efficient electrotransfection method for both mouse embryonic fibroblast (MEF) and induced pluripotent stem (iPS) cells based on the square-wave pulsing method using a Bio-Rad electroporation device. We developed the technology using different types of reporter plasmids and confirmed its very high efficiency for making gene knockout (KO) which was induced by both indels and targeted deletions using single or double gRNAs through CRISPR/Cas9 plasmids, respectively.

## Results

### Optimization of electrotransfection protocol for mouse iPS and EF cells

Three media were compared for electrotransfection efficiency. Initial results of transfection of mouse iPS cells with Bio-Rad electroporation buffer, PBS, and OptiMEM-GlutaMAX using a 10 ms single pulse of 200 V resulted in 84.7 % transfection rate in OptiMEM-GlutaMAX medium compared to 17.9 % in Bio-Rad buffer (Figure S1). More importantly, the number of cells and colonies 24 h after the electroporation was considerably higher in OptiMEM-GlutaMAX compared to both Bio-Rad and PBS buffers (p<0.05).

Then, we optimized pulse voltage, number, and duration as well as the medium temperature for electrotransfection of iPS cells based on the OptiMEM-GlutaMAX medium (Figures S2). We developed an electroporation protocol using the square-wave pulsing program of 250 V, 2 pulses, each 10 ms length, and 10 s interval in 4 mm cuvettes. Electroporations were carried out at room temperature with 20 µg DNA in 250 µl of OptiMEM-GlutaMAX. Applying this protocol for transient expression of mCherry and Venus reporters in mouse iPS cells showed a 95.5 and 92.8 % electrotransfection efficiency, respectively (Figure 1). Then, we implemented the same protocol with a higher voltage (300 V instead of 250 V) for MEF cells and reached 96.0 and 95.1 % expression of mCherry and Venus transgenes, respectively, 24 h after the electroporation (Figure 1). The electrotransfection efficiency was reproducible using mCherry reporter in three MEF and three iPS cell lines.

**Figure 1.**
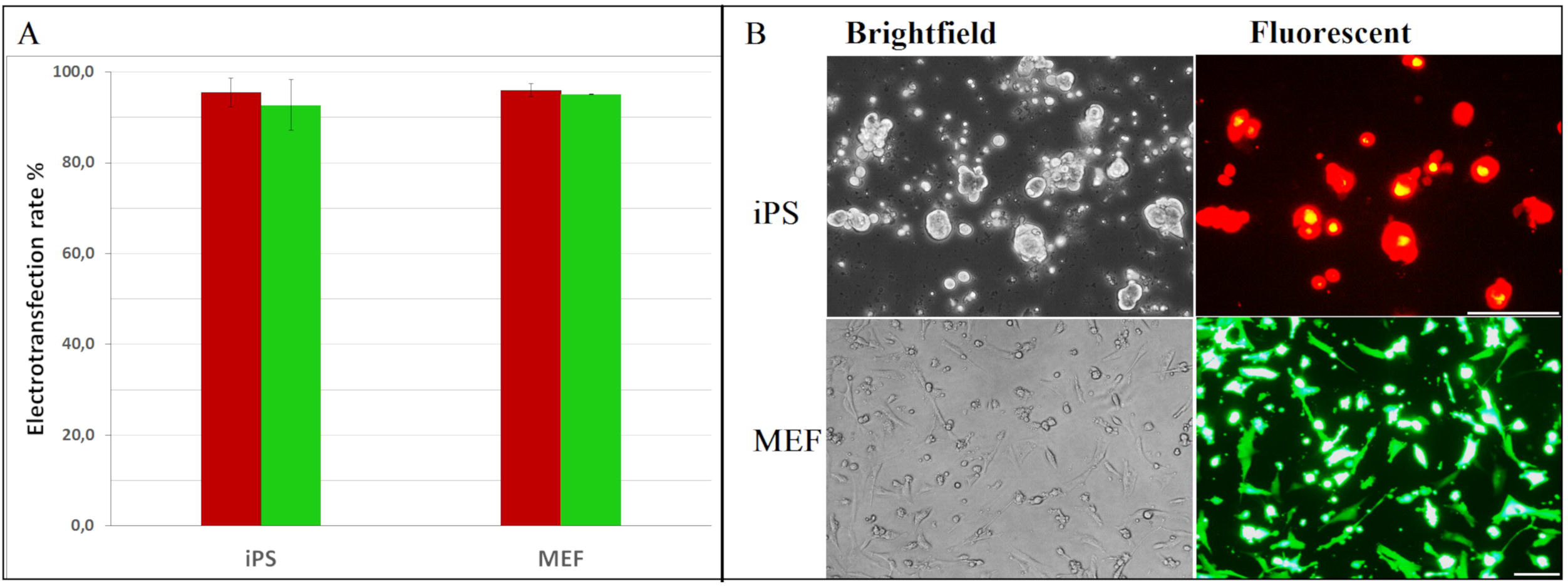
Electrotransfection efficiency of mouse embryonic fibroblast (EF) and iPS cells. In each electroporation reaction, 20 µg of either pT2-Venus and pT2-mCherry which encode Venus and mCherry proteins, respectively, was pre-mixed with cells and underwent electroporation. A) transfection efficiency and B) fluorescent expression of Venus and mCherry 36 h after electroporation. The following electroporation program was used: square-wave protocol with 300 V voltage, each 10 ms pulse length, 3 number of pulses, 10 s pulse interval, and 4 mm cuvette. Transfected cells for mCherry and Venus are depicted by red and green bars, respectively. Scale bars equal 100 µm. Results are means and standard deviation (SD).

Finally, using the optimized electrotransfection protocol the transient transfection efficiency of mouse iPS and EF cells was assessed in 250 µl of either of Bio-Rad, OptiMEM-GlutaMAX, and PBS media (Figure 2). Viability of iPS cells was significantly lower in Bio-Rad (22.0 %) and PBS (3.2 %) compared to the OptiMEM-GlutaMAX medium (78.2 %). Also, the electrotransfection efficiency of iPS cells was more than two-fold higher in OptiMEM-GlutaMAX (99.3 %) than that of Bio-Rad (42.7 %) and PBS media (47.2 %). The electrotransfection efficiency of mouse EF cells was 30 % higher in OptiMEM-GlutaMAX compared to the Bio-Rad medium (p-value <0.05). In addition, the viability of mouse EF cells averaged 78 % in the OptiMEM-GlutaMAX group.

**Figure 2.**
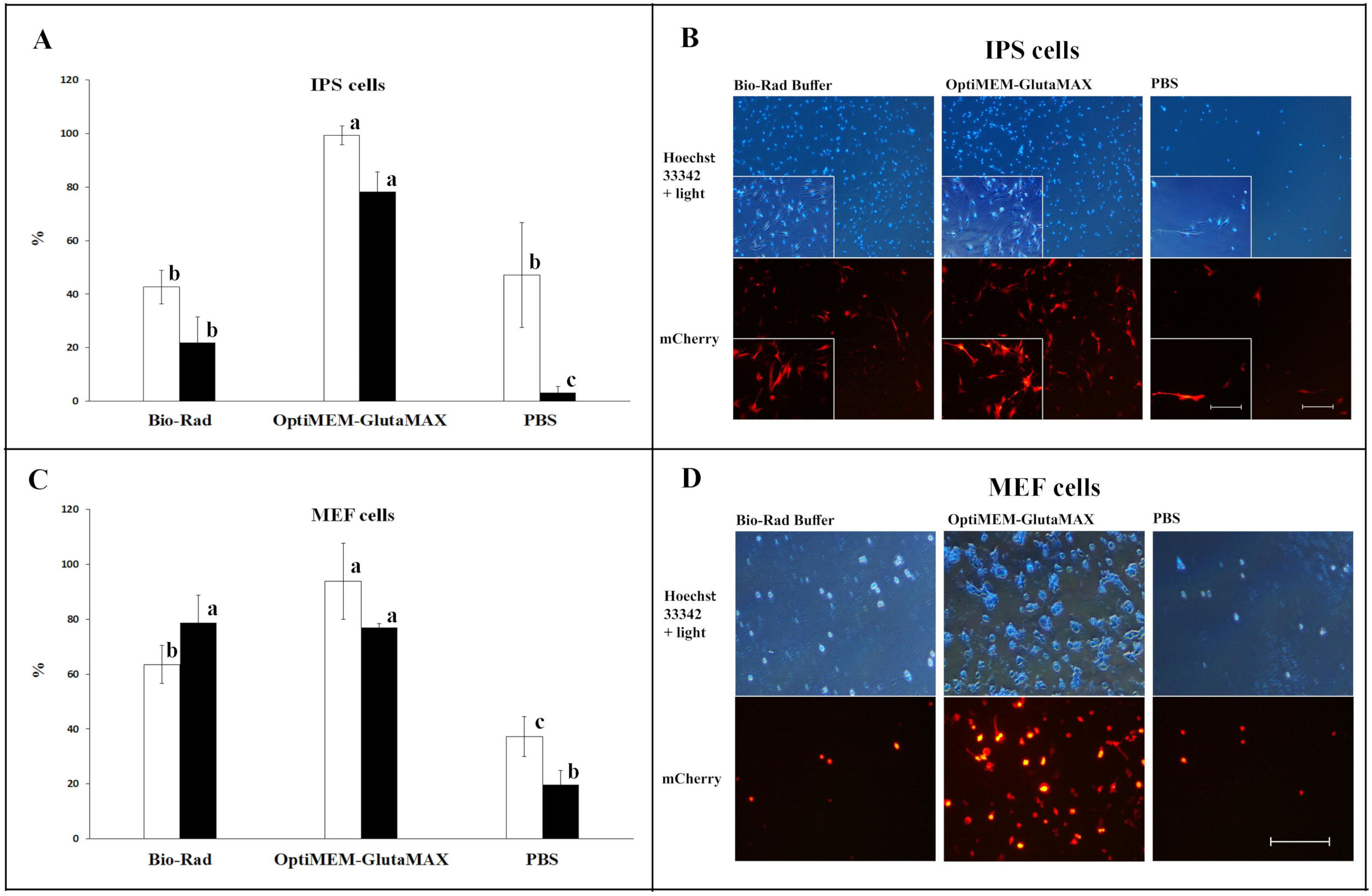
Cell electrotransfection and viability rates using Bio-Rad buffer, OptiMEM-GlutaMAX, and PBS. Electrotransfection efficiency of mouse iPS (A and B) and MEF cells (C and D) by BioRad electroporator. In each electroporation reaction, 20 µg of the reporter plasmid (encoding either Venus or mCherry) was pre-mixed with cells and underwent electroporation using the square-wave protocol with 250 and 300 V (for iPS and MEF cells, respectively), 2 pulses, each 10 ms length, 10 s interval, and 4 mm cuvette. The reporter expression was assessed 36 h after electroporation under a fluorescent microscope. White and black bars in A and C are electrotransfection efficiency and cell viability, respectively. Bars with different letters with are significantly different (p-value < 0.05). Scale bars equal 100 µm. Results are means and standard deviation (SD).

### Efficient knockout of Venus transgene using indels by Cas9/gRNA encoding plasmids

MEF cells carrying a single-copy of the Venus transgene were transfected by plasmids encoding a gRNA and Cas9 protein using the optimized electroporation method (Figure S3). Venus expression was not reduced by three gRNAs which targeted the upstream region of the Venus transgene although they induced indels in the targeted sites (Figure 3). However, targeting the beginning and ending parts of the transgene, ranging from +36 to +554 bp of the cDNA, could knockout the transgene in > 90 % of the electroporated cells (Figure S3). The Venus KO efficiency was maximized using gRNA+100 which was complimentary to the beginning part of the cDNA, so that only 2 % of cells maintained the functional Venus. The Venus knockout results were confirmed by fluorescent microscopy, as well as FACS analysis and DNA sequencing. The knockout efficiency of Venus transgene was not affected by the puromycin treatment; 99 % vs. 93 % for 100 gRNA with and without puromycin selection, respectively (Figure S4). Although electrotransfection results were very promising using OptiMEM-GlutaMAX medium, substituting the medium with the standard OptiMEM supplemented with glutamine was completely inefficient for making Venus KO (Figure S5). Moreover, electrotransfection of MEF cells with pSGD-Lys-72 plasmid (13.5 kb) encoding human lysozyme, puromycin, and gRNA-72 compared to the modified pX459-gRNA-72 showed a similar cell viability following a one-day selection against puromycin antibiotic (Figure S6).

**Figure 3.**
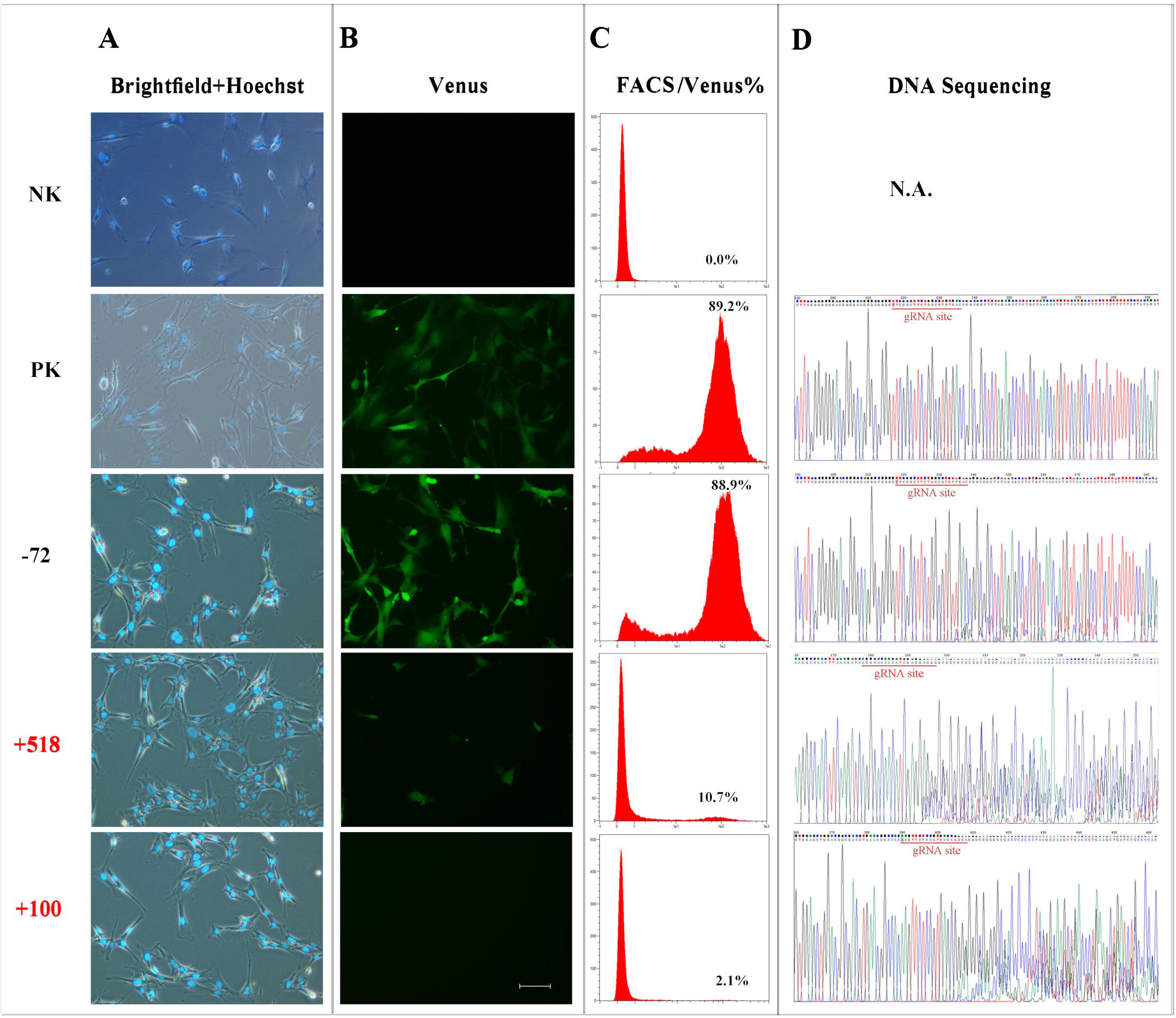
Knockout of Venus transgene by making indels using gRNA-encoding plasmids. MEF cells carrying a single-copy of the Venus transgene were electrotransfected with different gRNA-encoding plasmids (9.1 kb length). Induction of indels using gRNA-72 had no effect on the Venus expression, whereas gRNA+100 could completely remove the Venus signal. Electroporated cells were selected against puromycin and screened 10 days after the electroporation. Cells were stained with Hoechst 33342 and the efficiency of Venus knockout was assessed (A and B). Histoplots of the FACS results are shown with the percentage of Venus positive cells is embedded for each group (C). Results were confirmed by sequencing of the amplified Venus transgene (D). NK and PK are negative and positive controls, respectively. The following electroporation program was used: square-wave protocol with 300 V voltage, each 10 ms pulse length, 3 number of pulses, 10 s pulse interval, and 4 mm cuvette. Scale bar equals 100 µm.

### High efficient deletion of Venus fragments using dual gRNAs

Following making Venus knockouts using indels, we were interested in testing the efficiency of the electrotransfection protocol for making deletions inside of the target gene using two gRNAs. Simultaneously, MEF cells carrying a single-copy of Venus were co-electroporated using 15 combination of two gRNAs. The result of end-point PCR showed deletion of expected fragments ranging from 398 to 748 bp in the Venus transgene using all of gRNAs pairs (Figure 4). Only shortened Venus amplicons were amplified by a primer set (Figure S7), while using another primer set the original transgene with no deletion was also detectable (Figure 4). In overall, the deletion efficiency was more evident using the combinations of gRNA+676, although gRNA+554 and +518 were also efficient for making deletions. Results of the DNA sequencing confirmed the fragment deletion in co-transfected groups (Figure S8).

**Figure 4.**
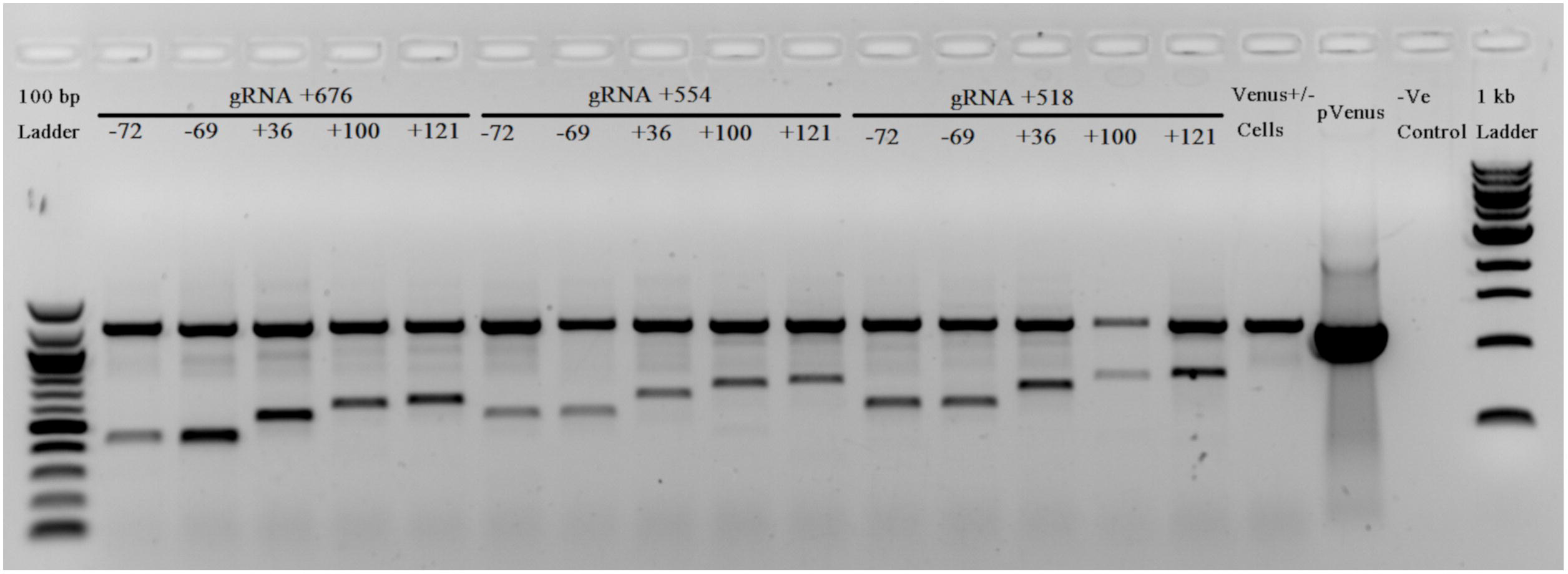
Targeted deletions of Venus transgene by co-electroporation of two gRNA-encoding plasmids. End-point PCR showed deletion of 15 different fragments from the original amplicon (1204 bp) using pairwise combination of gRNAs targeting the beginning (−72, −69, +36, +100, and +121) and the ending part (+518, +554, and +676) of Venus. Primer Venus Forward and Reverse1 amplified both the shortened and original fragments in the co-electrotransfected groups.

## Discussion

Nowadays, implementation of the nascent CRISPR/Cas9 technology is a straightforward approach to make genetic engineering in any organism. Electroporation of large plasmids (> 6 kb) is almost inefficient and associating with a low viability in primary cells (11). In the current study, we identified OptiMEM-GlutaMAX as the best medium for electroporation of MEF and iPS cells. The electrotransfection efficiencies were quite high throughout the initial setup condition with various pulsing conditions. A > 95 % electrotransfection rate was achieved with the optimized pulsing condition. Even the rate of transient expression of reporter genes, as well as the cell viability using the OptiMEM-GlutaMAX in the current study were as high as that of newly reported mechanical-electrical (7) and nanopore-electrical approaches (12). We used plasmids with various sizes, from 6.2 to 13.5 kb, and found a high transfection rate and cell viability irrespective of the plasmid size. It has been evident that electrotransfer of large plasmids (> 6 kb) associates with a very high cell toxicity, while the cell transfection rate was also low (11). However, in our study, electrotransfection of MEF cells using a 13.5 kb was as efficient as a 9.2 kb plasmid, whereas the cell viability remained the same. Therefore, the electrotransfection protocol for primary cells in the current study could overcome impediments of hard to transfect cells, as well as high toxicity, and low transfection rate using large plasmids (11). So far, invention of minicircles was considered as the main solution to overcome this problem (13). However, converting a plasmid to a minicircle is time- and cost-consuming and it does not solve the problem of co-transfection of plasmids (11). Nonetheless, our results showed that electrotransfection of large plasmids is possible with a high efficiency and cell viability in OptiMEM-GlutaMAX medium with no need for a long post-pulse recovery time, as suggested by Lesueur et al. (11). It has been well-documented that iPS cells are resistant and vulnerable to electroporation (14). The current study showed that comparing to the Bio-Rad electroporation buffer, the OptiMEM-GlutaMAX was compatible with iPS cells in terms of transfection rate and cell viability. However, unlike reported by Potter et al. (15), conducting the electroporation in ice-cold medium did not improve the transfection efficiency in the current study, although the transfection efficiency of ice-cold medium was almost independent of pulsing condition.

Then, we were interested to know if the transient expression is sufficient for genome engineering using the CRISPR/Cas9 technology. Following optimization of the electrotransfection protocol for reporter-encoding transgenes, we implemented the same protocol for transfer of plasmids encoding Cas9 and gRNA aiming to knockout the Venus transgene in MEF cells carrying a single-copy of Venus. The highest rate of Venus knockout through indels formation (> 95 % using gRNA+100) was as high as the electrotransfection efficiency of reporter-encoding plasmids. The indels rate using plasmids in the current study was higher than reported results by using both plasmid and RNP electrotransfection(16–18). Following making Venus knockouts using indels, we were interested to test the efficiency of the electrotransfection protocol for making deletions inside of the target gene using double gRNAs. Deletion of expected fragments in the Venus transgene by co-electroporation of two plasmids encoding two different gRNAs was confirmed using endpoint-PCR as well DNA sequencing. The deletion efficiency obtained in the current study is higher than previous results using dual gRNAs and Cas9 protein (19) as well as Cas9-encoding plasmids using the BTX electroporation buffer in mouse cell lines (20).

## Conclusion

In this study, we introduced a highly efficient electrotransfection method for both MEF and iPS cells based on the square-wave pulsing method using OptiMEM-GlutaMAX medium. This plasmid-based delivery method could induce > 95 % transfection efficiency for fluorescent reporter genes. Electrotransfection of Cas9/gRNA encoding plasmids caused in approximately 98 % knockouts of Venus transgene. Apart from indels creation, targeted deletions in the Venus transgene were achieved using the co-electroporation of two gRNA-encoding plasmids. Therefore, the developed protocol in the current study can be an alternate to using Cas9 protein and RNPs.

## Materials and methods

### Materials

All plastic consumables including cell culture flasks and plates, tubes, and filter tips have been purchased from SARSTEDT AG & Co. (Germany). Chemical reagents were purchased with the following information: Dulbecco’s phosphate buffered saline without calcium chloride and magnesium chloride (#D6662-10X1L, SIGMA-ALDRICH, Germany); OptiMEM 1X + GlutaMAX, reduced serum medium (#1854076, Gibco, Life Technologies, Germany); DMEM high glucose (4.5 g/l) w/o L-glutamine (#DMEM-HXA, Capricorn Scientific GmbH, Germany); L-glutamine for cell culture (#A3704,0100, Applichem, Germany); MEM nonessential amino acids solution (100X) (#NEAA-B, Capricorn Scientific GmbH, Germany); 2-mercaptoethanol (#BCBS5481, SIGMA-ALDRICH, Germany); penicillin/streptomycin solution (100x) (#PS-B, Capricorn Scientific GmbH, Germany); trypsin-EDTA (10X) (#L11-003, GE Healthcare, PPA Laboratories GmbH, Austria); fetal bovine serum (#10270-106, Gibco, ThermoFisherScientific, Germany); dimethyl sulfoxide (#D4540-500ML, SIGMA-ALDRICH, Germany); LIF (hBA-FL) (#sc-4377, Santa Cruz Biotechnology, Germany); sodium pyruvate (#P2256, SIGMA-ALDRICH, Germany); gelatin from bovine skin (#G939-100G, SIGMA-ALDRICH, Germany); and Hoechst 33342 (#62249, Thermo Scientific, Germany). The Gene Pulser Xcell™ system with CE Module from BIO-RAD (Germany) was used with 4 mm electroporation cuvettes (#748052, Biozym, Germany).

### Plasmids

#### Plasmids carrying *Sleeping Beauty* (SB) system

The *Sleeping Beauty* transposase was encoded in pCMV-T7_SB100X (4756 bp) which contains the hyperactive variant 100 of SB under the CMV promoter (21). Two reporter plasmids, pT2-Venus (6301 bp) encoding the Venus fluorescent marker under CAGGS promoter and pT2-mCherry (7756 bp) encoding mCherry, were used (21). In addition, a 13.5 kb plasmid, pSGD-Lys-72, was also used as a large plasmid which encodes human lysozyme under CAGGS promoter followed by Cas9 protein which was separated from puromycin resistance by a T2A peptide and expressed under CMV promoter, and gRNA-72 under U6 promoter. All pT2-Venus, pT2-mCherry, and pSGD-Lys-72 plasmids contained inverted terminal repeats (ITRs) of the SB transposase. The SB reprogramming transposon involved a CAGGS promoter driven cassette containing Oct4, Sox2, Klf4, and c-Myc, which were separated by sequences for coding the self-cleaving 2A peptides, and flanked by SB-ITRs (22).

#### Plasmid carrying the CRISPR/Cas9 system

For the CRISPR/Cas9 study, we modified pX459 plasmid (9151 bp) which encodes a Cas9 protein followed by puromycin resistance under CAGGS promoter/enhancer, as well as the gRNA scaffold under the U6 promoter. All gRNAs were synthetized and cloned into the backbone vector based on the Franham protocol (23). Briefly, gRNAs were selected to target Venus promoter or open reading frame (ORF) via CRISPOR software, available online (http://crispor.tefor.net/) (24). The list of gRNAs is available in Table 1. The 100 µM forward and reverse oligos were annealed in 10 µl reaction volume by incubation in a water bath containing pre-boiled water and letting it to cool down to the room temperature. The pX459 plasmid was digested with High Fidelity BbsI-HF (NEB #R3539) at 37°C for 10 min followed by gel purification using NucleoSpin Gel and PCR Clean-up Midi kit (#740986.20, MACHEREY/NAGEL). The purified fragment was kept at −20°C for further applications. Ligation of the annealed oligo-duplex with the digested pX459 was carried out as follow: diluted oligo-duplex (1:20 ratio from the 10 µM source) (1 µl), digested pX459 vector (50 ng), 10× T4 DNA ligase buffer (2 µl), and T4 DNA ligase (1 µl) in a 20 µl final reaction. The ligation reaction was incubated at room temperature for 1 hour. Transfection of the ligation mixture was carried out into NEB 5-alpha Competent E. coli (Cat. # C2987I) following incubation of 10 µl of the ligation reaction with the thawed competent cells on ice for 20 min and then at 37°C for 5 min. Then, 400 µl of SOC medium was added into the transformation tube, incubated at 37° C for 30 min, plated on Agar plates supplemented with 100 µg/ml ampicillin, and incubated at 37°C overnight. From the cultured plate, 10 colonies were picked and each was cultured in 3 ml LB medium followed by miniprep plasmid extraction (Genejet Plasmid miniPrep kit, #K0502). The extracted plasmids were simultaneously digested by BbsI and EcoRV FastDigest restriction enzymes (Fermentas). Plasmids with the expected bands of 6244 and 2928 bp were sent for sequencing.

### Cell culture and electroporation

#### Gelatinization of cell culture plates

A 0.1 % gelatin solution was prepared by stirring 250 ml PBS and 2.5 g gelatin for one hour without heating, and the solution was then autoclaved. Five milliliter of the 0.1 % gelatin was poured into a 6-well plate and aspirated after 1 min. Plates were left to be dried under the laminar flow hood for 30 min.

### Cell lines

We used mouse embryonic fibroblast cells (MEFs), carrying either no- or a single-copy of the Venus transgene (Venus +/-). Mouse embryonic fibroblast cells were isolated from day 11 embryos of the Venus −/− (wild type) or Venus +/− embryos. Embryos were beheaded, eviscerated, sliced, and cultivated with 5 ml fibroblast medium (DMEM with 10% FBS, L-glutamine, penicillin-streptomycin, nonessential amino acids, and β-mercaptoethanol) in T75 flasks. After 3 days of the incubation, propagated cells were trypsinized, passaged, and used for cell culture and transfection. Moreover, mouse iPS cells were derived from MEFs with no- or a single-copy of the Venus, respectively (22). Briefly, MEFs were electroporated with plasmids containing the 4-factor reprogramming transposon and the helper plasmid containing the hyperactive SB transposase (22). Electroporated cells were cultured in gelatinized 6-well plates and iPS medium. Presumptive iPS colonies were picked under microscope, and plated into individual wells of 96-well plates containing trypsin. Trypsin was neutralized with DMEM and 10% FCS, and the cells suspension was dispensed into individual wells of gelatinized 96-well plates (22).

### Cell Electroporation

To assess the transfection efficiency using reporter plasmids, MEFs and iPS cells with no-copy of Venus reporter were used. Both MEFs and iPS cells were cultured in 6-well plates. From the highly confluent culture, cells from each well of a 6-well plate were used for each electroporation reaction using both iPS and MEFs cells. Cells were washed once with 2 ml PBS and then trypsinized with 200 µl trypsin-EDTA. Trypsin activity was inhibited by 4 ml PBS per well, and then was removed through centrifugation at 1,000 rpm for 3 min. The cell pellet was resuspended into 250 µl of the electroporation medium. In an initial experiment, the highest transfection efficiency of iPS cells were reached by OptiMEM-GlutaMAX medium (Figure S1). Then, the effect of voltage and medium temperature was optimized for iPS cells (Figures S2 and S3). Finally, we reached the following optimized condition for electroporation of mouse iPS cells. Twenty micrograms of plasmid DNA were mixed with cells, and the cell-plasmid mixture was transferred into the 4 mm cuvettes, and underwent the following electroporation program: The square-wave protocol with 250 V, each 10 ms pulse length, 2 pulses, and 10 s pulse interval. MEFs underwent the same electroporation program which was optimized for iPS cells, except conduction of a 300 V. After the electroporation process, cells were transferred into the culture medium in 6-well plates and incubated at 37°C and 5 % CO_2_.

### Venus knockout using one and two gRNAs

For the CRISPR/Cas9 experiment, we used MEFs cells with a single-copy of Venus for making KO-Venus cells. The above-mentioned optimized protocol for cell electrotransfection was also used for Cas9/gRNA encoding plasmids. Knockout of Venus was carried out either by indels using 20 µg of a modified pX459 plasmid or by deletions via co-electroporation of two modified pX459 plasmids (each 20 µg per reaction). To consider the deletion efficiency of various sizes, three gRNAs which target the end part of Venus transgene (gRNA +518, +554, and +676) were pair-wisely co-electroporated with five gRNAs which targeted the beginning part of Venus (gRNA −72, −69, +36, +100, and +121). A pair of primers which cover all the gRNA complementary sites were used for the end-point PCR. DNA was extracted from each treated group using salting-out method. Briefly, confluent cells in each well of 6-well plates were trypsinized with 250 µl Trypsin-EDTA, directly transferred into a 1.5 ml tube containing 1 ml of cell lysis buffer (0.2 mg/ml proteinase K, 150 mM NaCl, 10 mM Tris, 10 mM EDTA, and 0.1% sodium dodecyl sulfate (SDS)) (25), and were incubated overnight at 55 °C in a shaker-incubator. Then, 500 µl saturated NaCl was added into each tube, converted for 5-10 times, and centrifuged with the high speed at room temperature. Supernatants were transferred into two new tubes and underwent ethanol precipitation with absolute ethanol followed by twice washing with 70 % ethanol using centrifugation with the high speed at 4°C. The DNA pellet was dissolved in 200 µl distilled water and kept at −20 °C till further usages.

### Fluorescent microscope imaging and flow cytometry analysis

For the fluorescent imaging, cells in each well of 6-wll plate were washed once with PBS, the medium was replaced with the transparent OptiMEM (without phenol) supplemented with Hoechst 33342 (2: 10,000 ratio from 1 mg/ml stock solution), and incubated at 37^°^C for 30 min. Transfection efficiency for mCherry and Venus reporters were assessed either by FACS machine or fluorescent microscopy. Cell viability was defined as cell numbers in the electroporated group divided by the cell number in the 0 voltage group (negative control). Knockout efficiency of various gRNA sequences targeting the Venus reporter was assessed 10 days after the electroporation. A flow cytometer, MACSQuant® Analyzer, was also used to assess cell transfection rate. We used Blue 488 nm in B1 channel with 525/550 nm filter and Yellow 461 nm Y2 channel with 615/20 nm filter for detection of Venus and mCherry proteins, respectively.

### Statistical analysis

Means comparison was carried out using the least significant difference (LSD) test (p-values 0.05). All experiments were carried out at least three times.

## Declarations

### Ethics approval and consent to participate

The German animal welfare law does not require an approval for humanely sacrifice of animals for the purpose of organ biopsies. The number of sacrified animals are annually reported to the regulating authorities.

### Consent for publication

Not applicable.

### Availability of data and materials

The datasets used and/or analysed during the current study are available from the corresponding author on reasonable request.

### Author’s contributions

Conceived the idea and designed the experiment: SE and WK; Carried out the experiments: SE, IH, and WK; Wrote the manuscript: SE; all authors read and confirmed the manuscript content.

## Acknowledgements

Authors would like to thank Antje Frenzel for conducting the FACS analysis.

