## Supplementary Tables and Figures for "A versatile bulk electrotransfection protocol for mouse embryonic fibroblast and iPS cells"

**Table S1.** The sequence of gRNAs specific for Venus transgene. These gRNAs were designed by CRISPOR online software.

| Name | gRNA sequence (5′-3′) |
| --- | --- |
| -252 | CTTCGCCCGCGCCCGCTAGA |
| -72 | TTCGGCTTCTGGCGTGTGAC |
| -69 | GGCTTCTGGCGTGTGACCGG |
| +36 | CGAGGAGCTGTTCACCGGCG |
| +100 | AAGTTCTCCGTGAGCGGCGA |
| +121 | GGCGAGGGCGACGCCACCTA |
| +518 | AGGCACAACATCGAGGACGG |
| +554 | AGCACGGGGCCGTCGCCGAT |
| +676 | CTGGAGTTCGTGACCGCCGC |

**Table S2.** Primers and probes for amplification and detection of Venus transgene

| Primer/probe Name | Sequence (5′-3′) | | Tm | |
| --- | --- | --- | --- | --- |
| Late Venus-Forward | | ACGCGTTAAGATACATTGATGAGTT | | 59.4 |
| Late Venus-Reverse | | CTTGTAGTTGCCGTCGTCCT | | 60.0 |
| Early Venus-Forward | | AGGACGACGGCAACTACAAG | | 60.0 |
| Early Venus-Reverse1 | | TGCCTTTTATGGTAATCGTGCG | | 59.6 |
| Early Venus-Reverse2 | | TTGCCTTTTATGGTAATCGTGCG | | 59.8 |
| Assay-Forward | | CCCTTCAGCTCGATTCTGTT | | 61.9 |
| Assay-Reverse | | GCAGGAGAGAACCATCTTCTT | | 61.7 |
| Probe | | 56-FAM-CCTCGGCTC-Zen-TGGTCCTGTAGTTGC-3IABkFQ | | 68.0 |

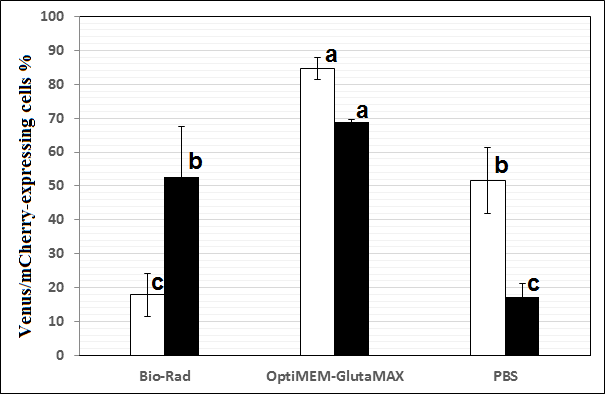

**Figure S1. Media effect on the electrotransfection efficiency of mouse iPS cells.** Three media (Bio-Rad buffer, OptiMEM-GlutaMAX, and PBS) were compared at the same electroporation condition at the early stage of protocol optimization. In each electroporation reaction, 20 µg of either pT2-Venus or pT2-mCherry was pre-mixed with cells and underwent electroporation using the square-wave protocol with 200 V voltage, 10 ms pulse length, and 4 mm cuvette. The expression of the fluorescent reporter was assessed 36 h after electroporation under a fluorescent microscope. White and black bars in A and C are electrotransfection efficiency and cell viability, respectively. Bars with different letters with are significantly different (p-value < 0.05). Results are means and standard deviation (SD).

mCherry-expressing cells%

| Pulse condition | Ice | RT |
| --- | --- | --- |
| 10 | 60.525 | 28.4 |
| 5_5 | 57.25 | 17.8 |
| 8_8 | 57.445 | 79.7 |
| 10_10 | 64 | 91.2 |

**Figure S2. The effect of temperature and pulsing condition on electroporation efficiency of mouse iPS cells**. Twenty microgram of a plasmid carrying mCherry transgene under CAGGS promoter was used for the electrotransfection. A square-wave protocol of 250 V, either single pulse for 10 ms or double pulses each for 5, 8, or 10 ms with 10 s pulse interval in 4 mm cuvettes, and 250 µl OptiMEM-GlutMAX was applied. Cuvettes were kept either on ice for 15 min (blue squares) or at room temperature (red triangles) before and after the pulsing conduction.

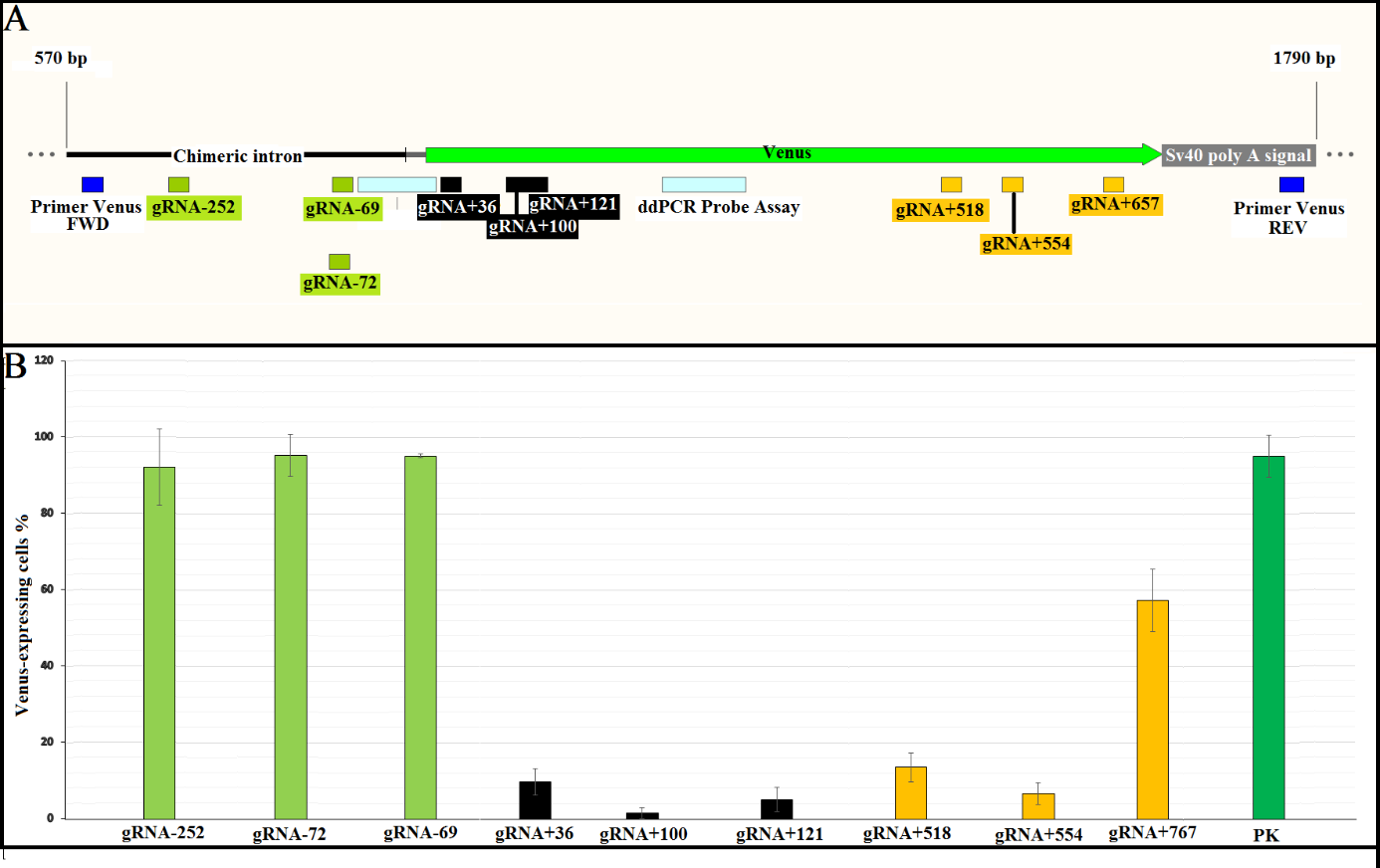

**Figure S3. Knockout of Venus transgene in MEF cells carrying a single-copy of Venus.** A) schematic presentation of gRNA location on the Venus transgene. Nine gRNAs were designed to target upstream region (252, 72, and 69), early encoding region (36, 100, and 121), and late encoding region of the Venus transgene (518, 554, and 676) which are depicted with white-green, black, and yellow bars, respectively. Assays for Digital PCR are depicted in white blue boxes. B) Efficiency of different gRNAs for making Venus knockout. Cells were treated with puromycin and were screened for the Venus transgene under a fluorescent microscope 10 days after the electroporation. PK is the positive control without handling which are MEF cells carrying a single-copy of the Venus transgene.

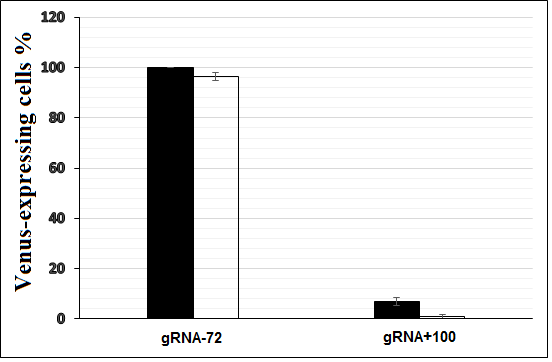

**Figure S4. Knockout efficiency of Venus transgene in MEF cells treated with or without puromycin.** Two plasmids encoding the Cas9 protein and gRNAs, 72 and 100, which targeted different positions of the transgene promoter and encoding gene were electroporated into the MEF cells carrying a single-copy of Venus using the optimized protocol. Cells were treated one-day post-transfection either with (white bars) or without Puromycin (black bars). Cells were screened for the Venus transgene under a fluorescent microscope 10 days after the electroporation.

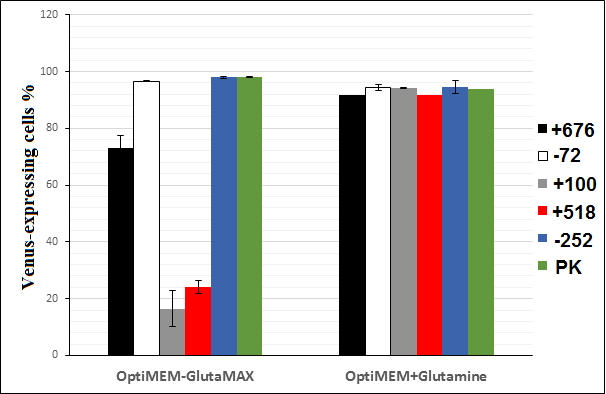

**Figure S5.** **Comparison of OptiMEM-GlutaMAX vs. standard OptiMEM (+Glutamine) for making Venus knockout.** MEF cells carrying a single-copy of Venus were electrotransfected using various gRNA-encoding plasmids (gRNA+676, -72, +100, +518, and -252). Cells were screened for the Venus transgene under a fluorescent microscope 10 days after the electroporation. PK is the positive control without handling which are MEF cells carrying a single-copy of the Venus transgene.

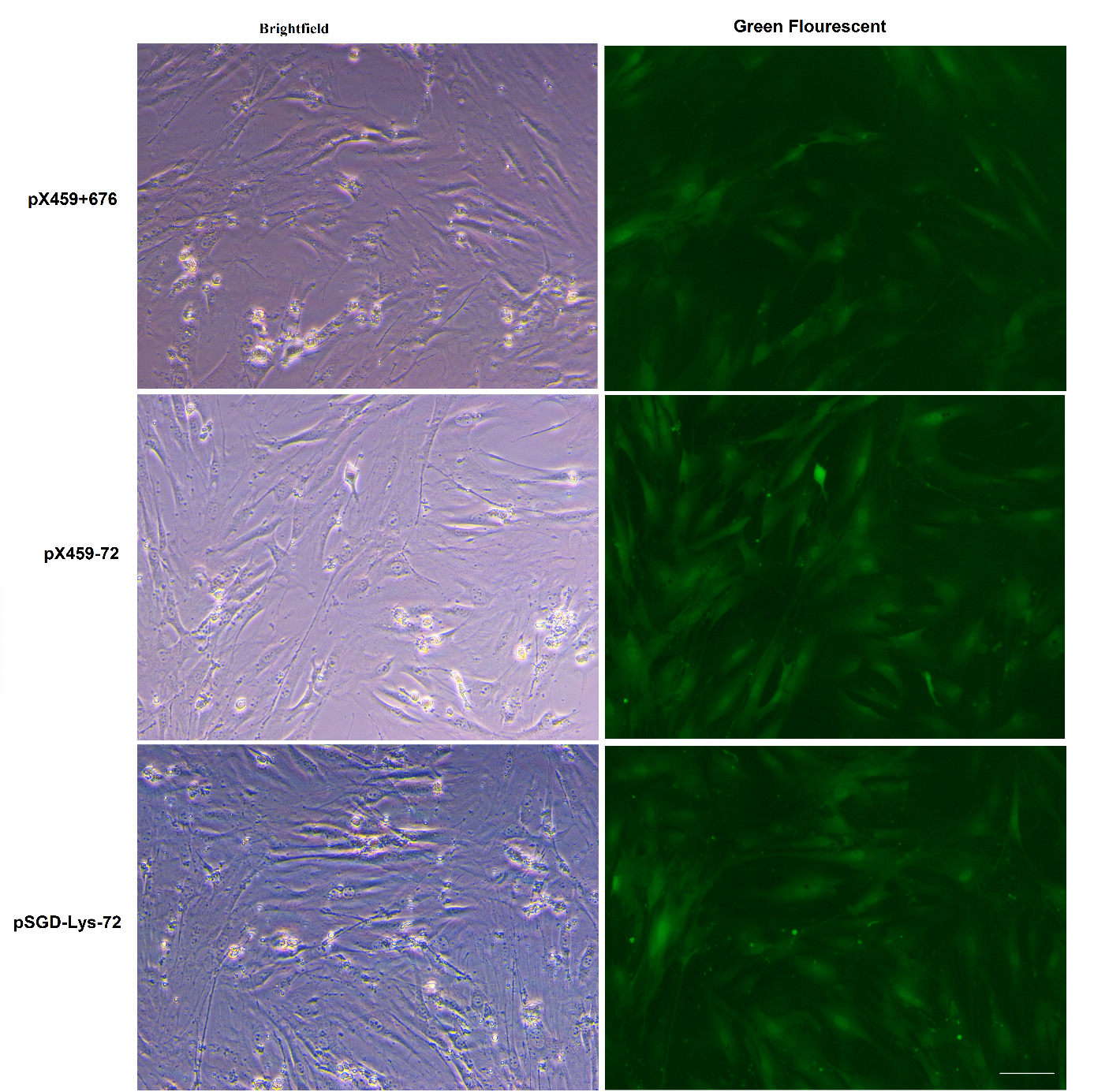

**Figure S6. Viability of electrotransfected cells using large plasmids.** Large plasmids ranging from 9.2 kb (pX459-72 and pX459+676) to 13.5 kb (pSGD-Lys-72) were used for electrotransfection of MEF cells carrying a single-copy of Venus. All plasmids encoded puromycin antibiotic. Cells from all groups underwent a one-day puromycin selection 16 h following the electroporation. The expression of Venus transgene was not affected by the electroporation during the first 40 h of cell culture. Scale bar equals 100 µm.

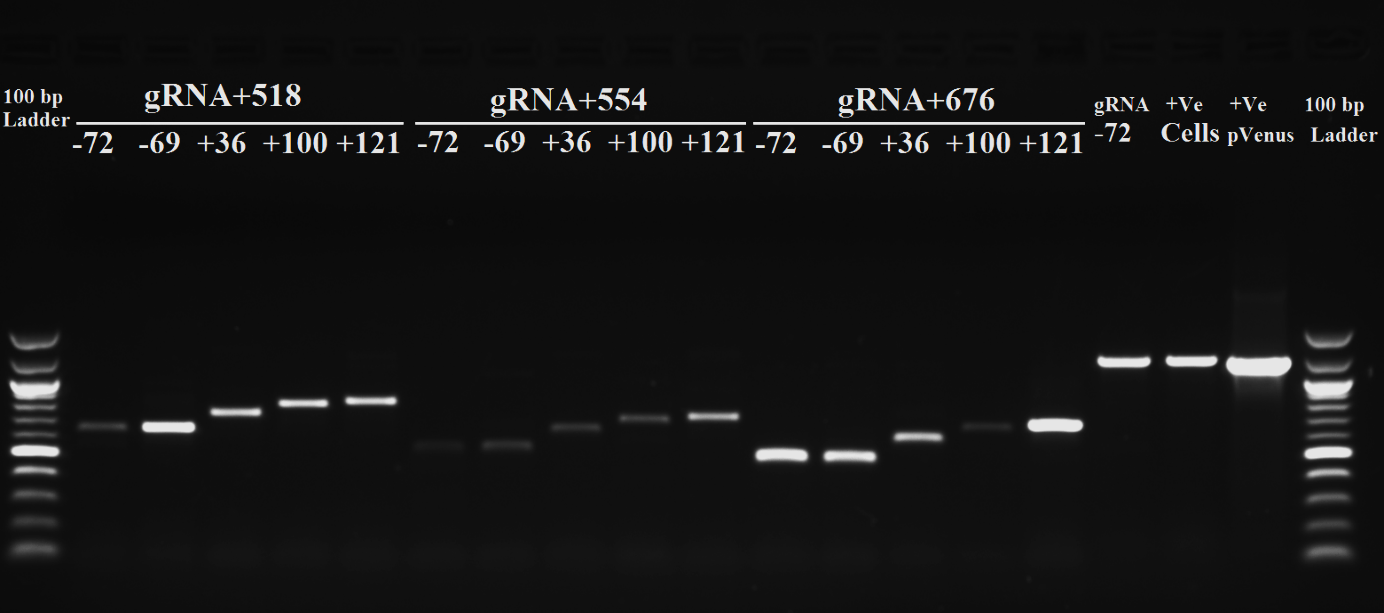

**Figure S7. Targeted deletions in Venus transgene by co-electroporation of two gRNA-encoding plasmids.** End-point PCR using primer Venus Forward and Venus Reverse showed deletion of five different fragments from the original amplicon (1204 bp) using pairwise combination of gRNAs targeting early (-72, -69, +36, +100, and +121) and late part (+518, +554, and +676) of Venus. Only the shortened fragments have been amplified using this primer set in the co-electrotransfected group.

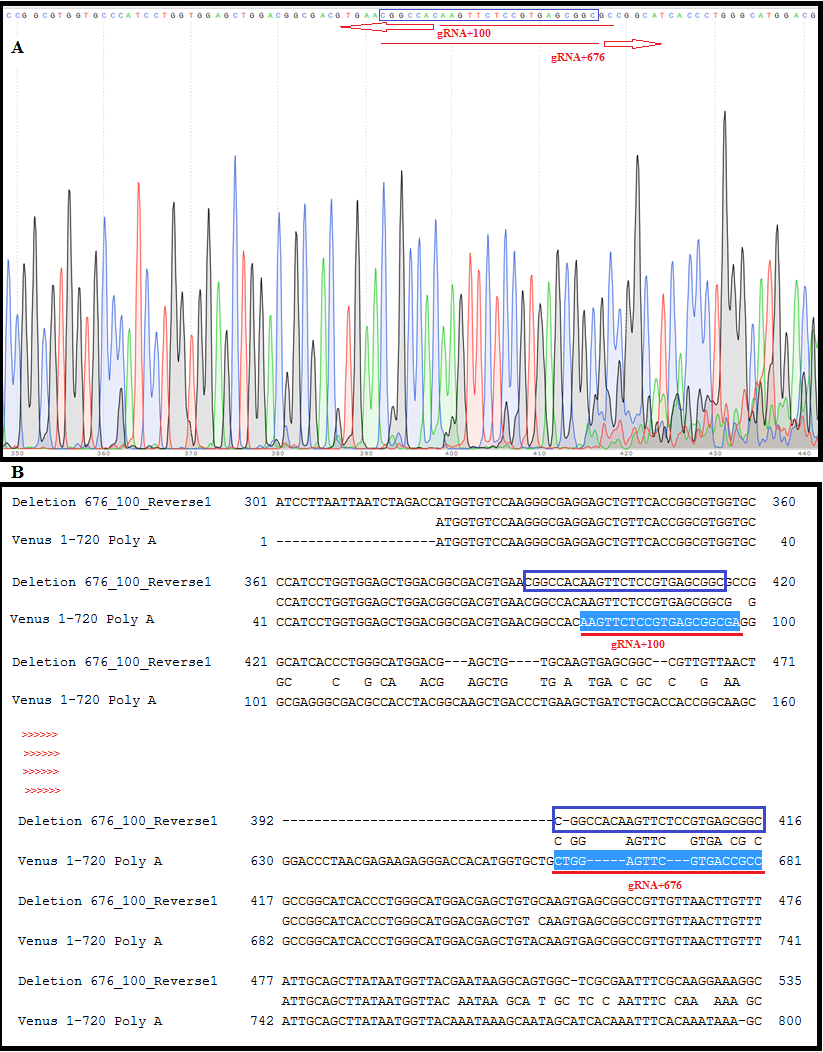
**Figure S8. Sequencing results of Venus transgene which underwent a targeted deletion.** MEF cells carrying a single-copy of Venus were co-electrotransfected with two gRNA-encoding plasmids. A) The amplified DNA from cells which were treated with gRNA+100 and gRNA+676 were sequenced. B) Alignment of the sequenced DNA with the Venus sequence. Blue box indicates a common sequence which corresponded to both gRNAs. The gRNAs and their following sequences are depicted with red color.
